# Microstimulation of human somatosensory cortex evokes task-dependent, spatially patterned responses in motor cortex

**DOI:** 10.1101/2022.08.10.503543

**Authors:** Natalya D. Shelchkova, John E. Downey, Charles M. Greenspon, Elizaveta V. Okorokova, Anton R. Sobinov, Ceci Verbaarschot, Qinpu He, Caleb Sponheim, Ariana F. Tortolani, Dalton D. Moore, Matthew T. Kaufman, Ray C. Lee, David Satzer, Jorge Gonzalez-Martinez, Peter C. Warnke, Lee E. Miller, Michael L. Boninger, Robert A. Gaunt, Jennifer L. Collinger, Nicholas G. Hatsopoulos, Sliman J. Bensmaia

## Abstract

Motor (M1) and somatosensory (S1) cortex play a critical role in motor control but the nature of the signaling between these structures is not known. To fill this gap, we recorded – in three human participants whose hands were paralyzed as a result of a spinal cord injury – the responses evoked in the hand and arm representations of primary motor cortex (M1) while we delivered ICMS to the somatosensory cortex (S1). We found that ICMS of S1 activated some M1 neurons at short, fixed latencies, locked to each pulse in a manner consistent with monosynaptic activation. However, most of the changes in M1 firing rates were much more variable in time, suggesting a more indirect effect of the stimulation. The spatial pattern of M1 activation varied systematically depending on the stimulating electrode: S1 electrodes that elicited percepts at a given hand location tended to activate M1 neurons with movement fields at the same location. However, the indirect effects of S1 ICMS on M1 were strongly context dependent, such that the magnitude and even sign relative to baseline varied across tasks. We tested the implications of these effects for brain-control of a virtual hand, in which ICMS was used to convey tactile feedback about object interactions. While ICMS-evoked activation of M1 disrupted decoder performance, this disruption could be minimized with biomimetic stimulation, which emphasizes contact transients at the onset and offset of grasp, reduces sustained stimulation, and has been shown to convey useful contact-related information.

**Significance:** Motor (M1) and somatosensory (S1) cortex play a critical role in motor control but the nature of the signaling between these structures is not known. To fill this gap, we recorded from M1 while delivering intracortical microstimulation (ICMS) to S1 of three human participants, whose hands were paralyzed by spinal cord injury. We found that ICMS activates M1 and that the motor fields of activated M1 neurons match the sensory fields of the stimulated S1 electrodes. These findings have important implications for using ICMS to convey tactile feedback for brain-controlled bionic hands. Indeed, the ICMS-evoked M1 activity worsens control of the hand. Fortunately, this effect is minimized by using biomimetic tactile feedback, which emphasizes contact transients and reduces sustained ICMS.

## Introduction

Manual interactions with objects involve the integration of sensory signals – about the state of the hand and its interactions with objects – and motor signals – about intended actions. Dexterous hand use relies on both somatosensory and motor cortices as evidenced by the severe deficits in manual dexterity that follow lesions to either of these brain regions^1,2^. However, many of the cortical mechanisms of sensorimotor integration remain to be elucidated. Brodmann’s area 1 of somatosensory cortex (S1) has been shown to send projections, albeit sparse ones, to primary motor cortex (M1)^3,4^, and this direct sensorimotor pathway has been hypothesized to play a key role in integrating sensory signals into motor execution. Electrical stimulation of human S1 has been shown to evoke responses in the local field potentials measured in M1^5,6^, consistent with the identified anatomical pathway. However, the modulation of single-cell responses in M1 to S1 stimulation and the function of the signals passed from S1 to M1 remain largely unknown.

To fill this gap, we delivered – in three human participants whose hands were paralyzed as a result of a spinal cord injury – intracortical microstimulation (ICMS) to the hand representation of S1 while we recorded the responses in the hand and arm representation of M1. First, we quantified the prevalence and temporal characteristics of ICMS-evoked activation. Second, we characterized the spatial pattern of activation in M1 and its relationship to the location of the stimulating electrode. Third, we compared ICMS-evoked M1 activity in different task conditions. Finally, we assessed the consequence of the ICMS-evoked activity on our ability to infer motor intent from M1 signals.

## Results

ICMS pulse trains varying in frequency and amplitude were delivered under two conditions: a passive condition in which the participants watched videos and an active condition in which the participants attempted to reach toward, grasp, and transport a virtual object, a task commonly used for BCI calibration^7^.

### Motor cortex responds to stimulation of somatosensory cortex

First, we examined the responses of M1 neurons to a 60-μA, 100-Hz, 1 second duration ICMS pulse train delivered through individual electrodes in S1 in the passive condition (see Figure 1A and Supplementary Figure 1 for array locations). We found that, on a large subset of electrodes in M1, ICMS of S1 modulated the activity of multi-units in M1 (Figure 1B and Figure 2). In some cases, ICMS had an excitatory effect on M1, leading to an increase in activity, (Figure 1B, left) and in others it had an inhibitory effect, leading to a decrease (Figure 1B, right). We verified that these effects were not electrical artifacts by confirming that they were also observed in the responses of sorted single units (Supplementary Figure 2 and Supplementary Figure 3). The prevalence and strength of these effects varied across participants: participant C1 showed more prevalent and stronger effects than the other two (P2 and P3, Figure 2, Supplementary Figure 4). The participants also differed in the sign of the ICMS-induced modulation, with primarily excitatory responses in C1 (94.2%) and a more even mix in P2 and P3 (39.3% and 46.0% excitatory, respectively).

**Figure 1.**
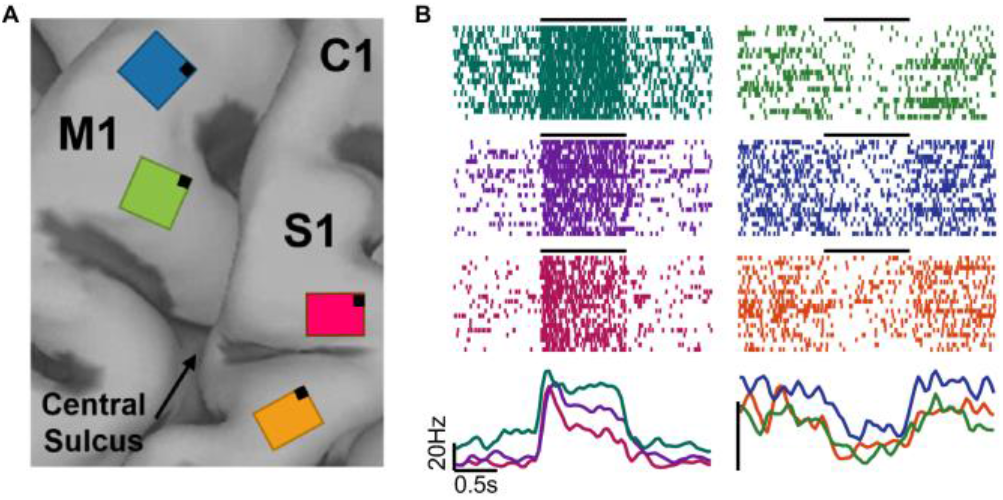
A| Four Neuroport electrode arrays (Blackrock Microsystems, Inc.) were implanted in the hand and arm representations of motor cortex (M1) and the hand representation of somatosensory cortex (S1). Here, the implant locations are shown for participant C1. Black squares on the arrays indicate the posterior-medial corner of each array, which is used as a reference in later figures. B| M1 responses to ICMS trains delivered to S1. Responses of three example motor channels (spike rasters above and averaged, smoothed firing rates below) that were excited by ICMS (left) and three that were inhibited by ICMS (right). Black horizontal lines indicate ICMS period. Top two from participant P3, bottom four from participant C1.

**Figure 2.**
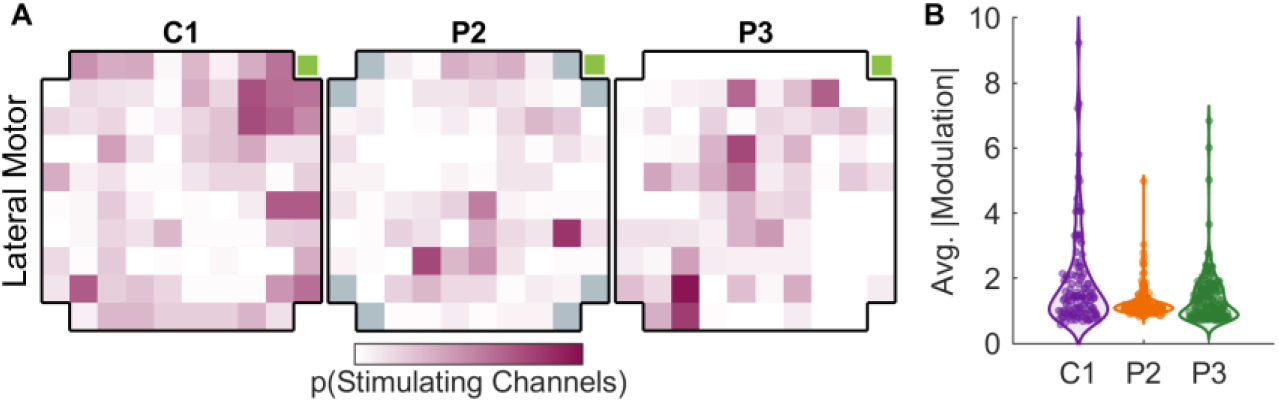
Prevalence of ICMS-evoked activity in motor cortex. A| Proportion of stimulating channels that significantly modulated each motor channel on the lateral motor array of each participant (range: 0 - 0.7). In P2, grey squares indicate channels that are not wired. The majority of motor channels could be modulated by ICMS through at least one channel in S1. The green square indicates the posterior-medial corner of the array (see Figure 1A and Supplementary Figure 1). B| ICMS-driven modulation of activity in each M1 channel, averaged across stimulating channels. Modulation is the ICMS-driven change in the response, normalized by baseline fluctuations.

### Stimulation of somatosensory cortex can directly activate neurons in motor cortex

Next, we examined whether the ICMS-evoked activation of M1 was driven by direct input from S1, reflected in responses that are temporally locked to the stimulation pulses. To this end, we computed the pulse-triggered average for each pair of stimulating and recording electrodes. We found M1 channels with responses that were systematically locked to the stimulation pulses (Figure 3A; Supplementary Figure 5). For most of these channels, the evoked neural activity occurred between 2 and 6 ms after pulse onset with millisecond or even sub-millisecond jitter across pulses (Figure 3B; Supplementary Figure 6). To eliminate the possibility that the response latency was longer than the inter-pulse duration, we measured the latency with pulse trains at different frequencies (25, 50, and 100 Hz) and found the latency to be consistent across frequencies (Supplementary Figure 7). Of the motor channels that were modulated by ICMS delivered to S1, 37.3%, 0.6%, and 31.8% exhibited this pulse-locked response in C1, P2, and P3, respectively. Most channels showed large and significant shifts in firing rate during bouts of ICMS with no pronounced peak in the pulse-triggered average (Figure 3C; Supplementary Figure 8). Thus, some of the ICMS-evoked activity in M1 seems to be triggered through direct activation from S1, possibly monosynaptically, whereas the majority seems to reflect more indirect effects.

**Figure 3.**
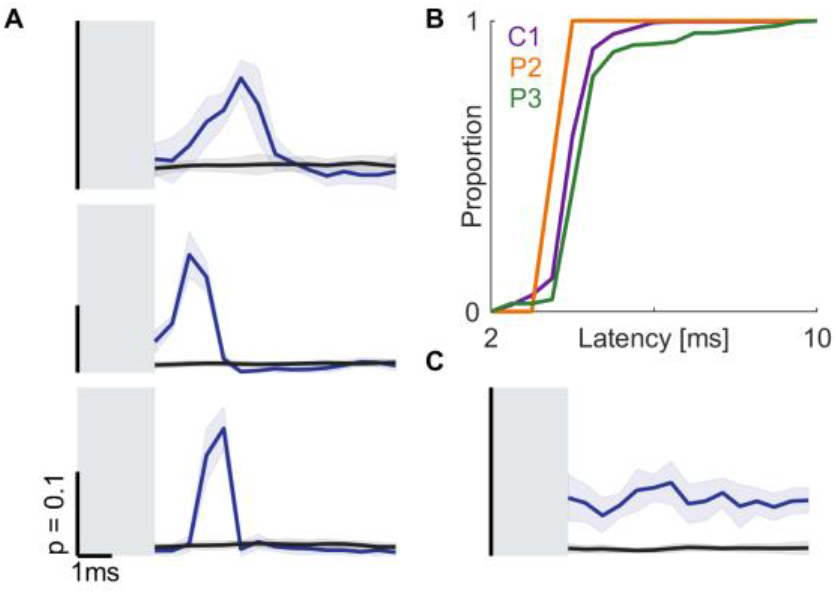
Short-latency, pulse-locked responses in M1. A| Pulse-triggered average of the responses of three motor channels to 100-Hz ICMS. On a subset of channels, such as these, responses were tightly locked to each pulse with millisecond or even sub-millisecond jitter across pulses. Blue line is during stimulation, black is during baseline (sham stimulation), grey box indicates blanked recording time to eliminate the stimulation artifact. Scale bar indicates a 10% probability of a spike occurring in a 0.5-ms bin. B| Cumulative distribution of the latency of the peak pulse-locked response. Latencies tended to be shorter than 6 ms. C| Pulse-triggered average of the response of a motor channel whose activity increases with stimulation but is not pulse locked.

### Spatial pattern of activation in motor cortex varies systematically across stimulating electrodes

Next, we examined the spatial pattern of ICMS-evoked activity over the M1 surface (both direct and indirect) and assessed whether the activated M1 channels differed systematically across stimulating electrodes. We found that different stimulating electrodes evoked different spatial patterns of activation in M1 (Figure 4A). Moreover, these patterns changed systematically: neighboring stimulating electrodes tended to produce similar patterns of M1 activation whereas distant stimulating electrodes evoked different, sometimes non-overlapping activation patterns. This was particularly pronounced when comparing the spatial pattern of M1 activation across electrodes in different sensory arrays (Figure 4B, Supplementary Figure 9) but was also observed when comparing activation across electrodes in the same sensory array (Supplementary Figure 10). While the trends were similar, P2 showed lower correlations in ICMS-evoked spatial patterns and a smaller (though still significant) difference between the within-array and cross-array correlations.

**Figure 4.**
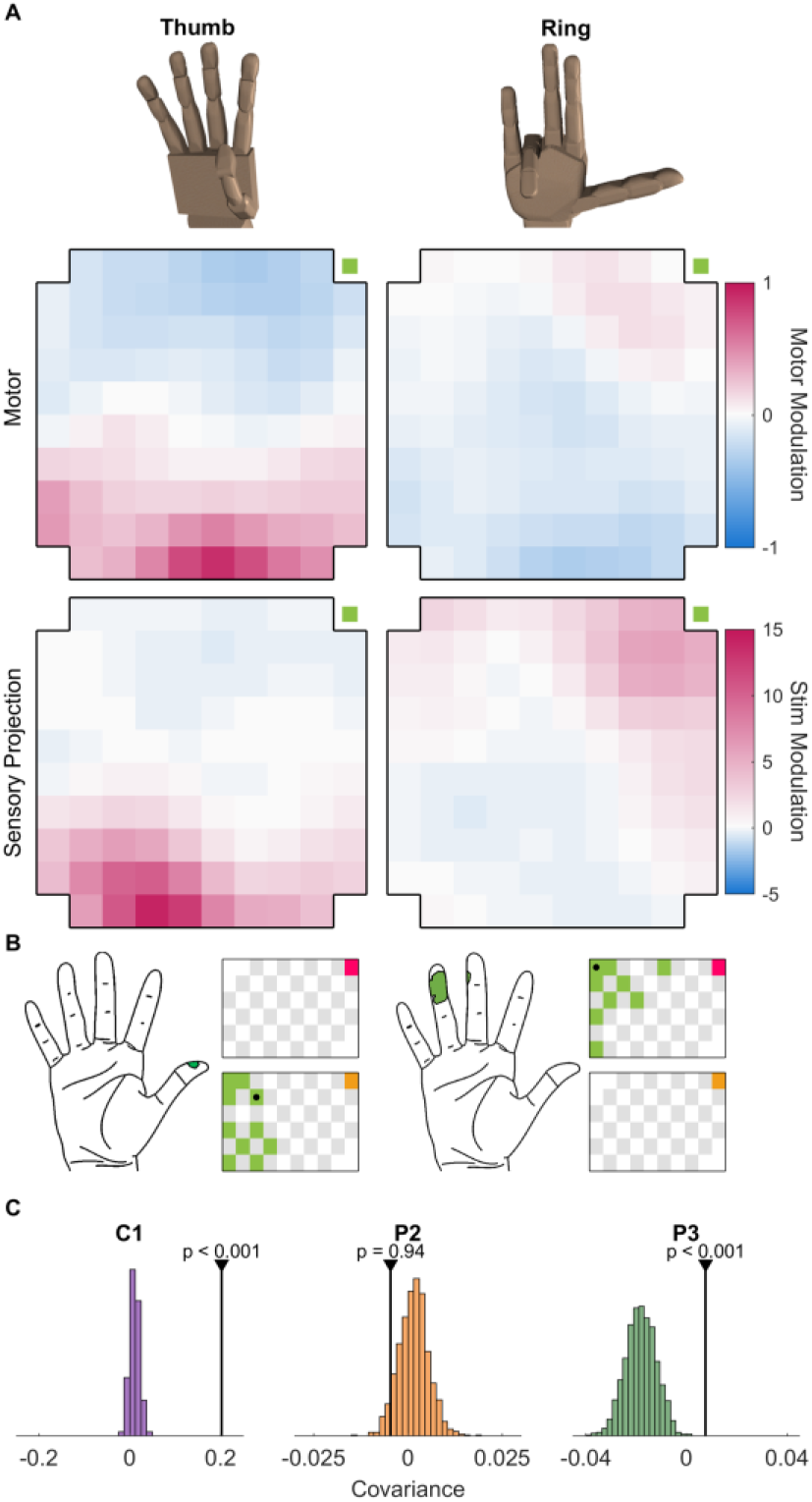
Shared somatotopy between movement-evoked and ICMS-evoked activity in participant C1. A| Top: Rendering of the extrema of thumb and ring flexion in virtual reality. Bottom: Difference in activation during attempted flexion of the thumb (left) and ring finger (right) vs. the mean activation during attempted flexion of each of the 5 digits. B| Top: Average M1 activity evoked by stimulation through S1 channels with projected fields on the thumb (left) and the ring finger (right). Green squares indicate the posterior and medial corner of the array (Figure 1A). Bottom: Projected fields reported by participant C1 when stimulated through one channel in the lateral and medial sensory array, respectively (indicated by a black dot in the array maps). Channels shaded in green denote electrodes with projected fields on the thumb and ring finger, respectively. Channels shaded in gray denote unwired electrodes. Pink and orange squares in the top right indicate the posterior and medial corner of the medial and lateral sensory array, respectively (Figure 1A). Motor channels that respond strongly to attempted thumb or ring finger movements tend to also be strongly activated by stimulation of electrodes with projected fields on the thumb or ring finger, respectively. C| Measured covariance between motor and sensory projection maps for the lateral motor arrays in C1, P2, and P3 along with the null distribution.

Examination of the spatial patterns of M1 activation suggests a coordinated progression. In participant C1, for example, lateral stimulating electrodes tend to activate neurons on the lateral aspect of M1 and medial stimulating electrodes tend to activate neurons on the medial aspect (Figure 4A). We hypothesized that this progression reflects the respective somatotopic organizations of S1 and M1. That is, stimulation through electrodes with projected fields on the thumb - those that evoke a sensation experienced on the thumb during stimulation - might preferentially activate M1 neurons that drive thumb movements. To test this hypothesis, we mapped the somatotopic organization of M1 by measuring, on each motor channel, the evoked activity when the participant attempted to move each digit. For each motor channel, we computed the difference between the activation evoked during attempted movement of each digit and the mean activation during movement of each of the five digits (motor map, Figure 5A). We mapped the somatotopic organization of S1 by identifying the digit on which the participant reported the sensation when stimulation was delivered through each electrode (projected field, bottom of Figure 5B). Having constructed these motor and sensory maps, we then derived the pattern of M1 activation when ICMS was delivered through electrodes with projected fields on each digit in turn (sensory projection map, Figure 5B). Finally, we assessed the degree to which the motor map matched the sensory projection map. For example, we asked: to what extent are M1 electrodes that maximally respond during attempted thumb movements activated when stimulation is delivered through S1 electrodes with projected fields on the thumb? To answer this question, we computed the covariance between the motor and sensory projection maps and compared this covariance with that obtained when the maps were scrambled. We found the match between motor and sensory projection maps to be far greater than that expected by chance (p < 0.001, permutation test) for participants C1 and P3 (Figure 5C), consistent with the hypothesis that electrical activation of S1 neurons leads to activation of M1 neurons with matching movement fields (Supplementary Figure 11).

**Figure 5.**
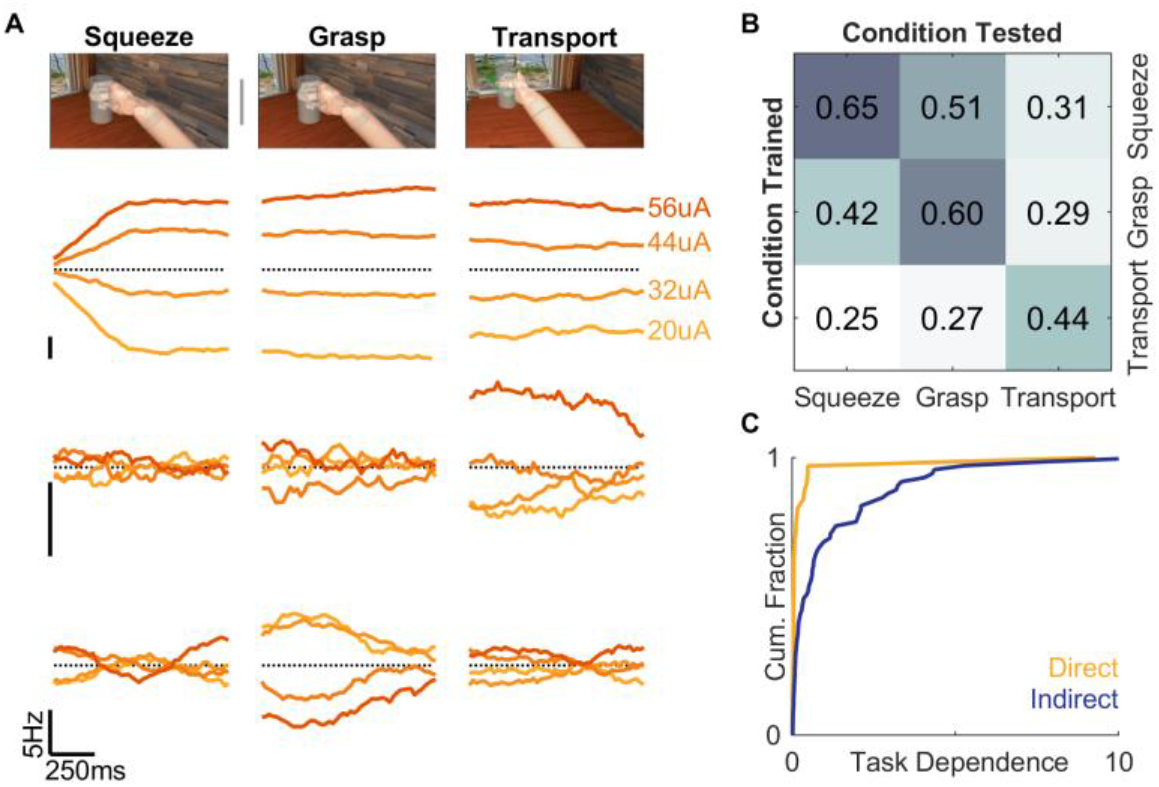
ICMS-evoked activity depends on behavior. A| Top: Squeeze, grasp, and transport in the VR environment. Bottom: Three example motor channels from Participant C1 exhibit different responses to four levels of ICMS across three motor conditions (squeeze, grasp, transport). Traces represent the mean subtracted firing rate evoked by stimulation at the four levels. The vertical and horizontal scale bars denote a firing rate of 5 Hz and a timespan of 250 ms, respectively. B| Stimulation amplitude classifier performance. Classifiers were trained from M1 activity on one of the three conditions and tested on activity in each condition (cross-validated within condition). C| Task dependence – gauged by the strength of the condition/amplitude interaction divided by the strength of the main effect of amplitude – is nearly zero for the pulse-locked responses (direct) but varies widely for the non-pulse locked (indirect) ones. The response of units with direct input from somatosensory cortex respond the same way to ICMS across behavioral conditions. Kolmogorov-Smirnov test; Figure 7B; Supplementary Figure 13). In other words, the ICMS contaminated the neuronal activity used by the decoder to infer motor inte nt. The disruption is likely to be far stronger when decoding hand (rather than arm) movements given that ICMS-driven activity in M1 is strongest for somatotopically linked segments.

In contrast, the somatotopic patterning was much weaker and non-significant in participant P2 (Supplementary Figure 11). Note, however, that ICMS-driven M1 activation in this participant was sparse (Figure 2A), weak (Figure 2B), and unpatterned (Supplementary Figure 9C,D). We hypothesize that the lack of spatial patterning reflects the fact that this participant’s most lateral M1 array was more medial than were its counterparts in the other two participants, and thus likely located in the proximal limb representation rather than hand representation. Consistent with this hypothesis, there was very little digit-specific activity in the M1 array of P2 during the individuated digit movement task (Supplementary Figure 11B). The motor arrays in participant P2 are much older than are those in participants C1 and P3, which may have contributed to the observed differences.

### Stimulation-evoked activation in motor cortex differs across tasks

The analyses shown above were carried out on M1 responses collected when the participants were not engaged in any motor task. Next, we examined whether ICMS-evoked M1 activity was task-dependent. To this end, we measured ICMS-evoked responses in M1 as participants C1 and P3 performed two tasks. In the first task (‘squeeze’), they attempted to squeeze a cylinder in a virtual reality environment (without making any overt movement). In this task, contact with the virtual cylinder triggered ICMS (frequency = 100 Hz) through two electrodes delivered at one of four amplitudes (20, 32, 44, and 56 μA, presented in random order). The participants were instructed to report the magnitude of the ICMS train to ensure their engagement. In the second task (grasp and transport), the participants observed and attempted to mimic the actions of a virtual limb as it reached for and grasped the cylinder in one location and transported it to a different location. Upon grasp, the same ICMS trains were delivered as in the squeeze task (again ordered randomly across trials) until the object reached the target location. For this task, we analyzed the responses during the grasp phase and the transport phase separately. We reasoned that the grasp epoch involved the same behavior as did the ‘squeeze’ task, whereas the transport phase involved a different behavior. We then compared M1 responses to ICMS across the three conditions (‘squeeze,’ ‘grasp,’ and ‘transport’).

We first verified that M1 was engaged in the two behavioral tasks by examining the task dependence of the M1 activity. We found that activity on most motor channels differed across task conditions (squeeze vs. grasp vs. transport, >80% of the electrodes exhibited significant task modulation according to a multi-way ANOVA, p < 0.05 in both participants). Moreover, the observed reach endpoint could be decoded during the grasp and transport task from the M1 population activity (84 and 87% classification accuracy for two sessions with participant C1 and 26% accuracy for participant P3; chance = 12.5%; in participant P3, the motor arrays were much more strongly modulated by hand/wrist than shoulder movements, thus the poor performance).

Examining the dependence of the M1 activity on ICMS amplitude, we found that a large number of motor channels were modulated in an amplitude dependent way (across participants and pairs of stimulating electrodes, 31% and 78% in two different sessions with participant C1; 54% in participant P3; p < 0.05 multi-way ANOVA). Surprisingly, however, the effect of ICMS varied significantly across tasks for a large number of units (across participants, 17%, 42%, and 19% of the modulated channels exhibited significant task-amplitude interaction for the two sessions with participant C1 and that with participant P3, p < 0.05) (Figure 6A, Supplementary Figure 12A). For example, the responses of some neurons in M1 were strongly modulated by ICMS during some tasks but not others. Even the ‘squeeze’ and ‘grasp’ conditions sometimes yielded different ICMS-evoked modulation, even though the behavior is nearly identical – the only difference being that grasp occurs at the end of a reach and just before transport whereas squeeze does not. To further demonstrate the task dependence of the ICMS effects, we built a classifier of ICMS amplitude based on responses obtained in one of the three conditions (squeeze, grasp, transport) and attempted to use it to decode ICMS amplitude from the responses in the other two conditions. We found that, while we could decode ICMS amplitude on held-out data within condition with up to 65% accuracy, performance was worse across conditions (Figure 6B, Supplementary Figure 12B). In particular, the effects of ICMS in the transport condition were very different from those in the squeeze or grasp conditions as evidenced by the poor performance of classifiers built on one task and tested on the other.

**Figure 6.**
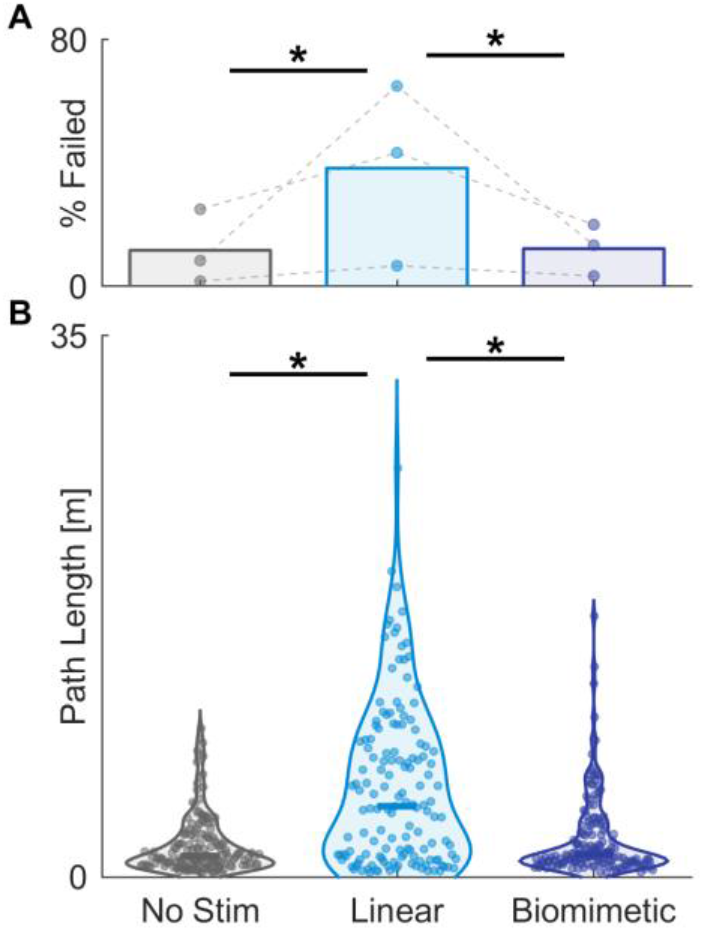
Decoder performance with and without sensory feedback (from participant C1). A| Failure rate for the three conditions. Rates collected during a single session are connected by a dotted line. B| Path length during the transport phase with different stimulation conditions. Linear stimulation caused the path length of the transport phase to be significantly longer than without stimulation (p < 0.001, K-S Test). In contrast, biomimetic stimulation was significantly more efficient than its linear counterpart and not significantly different from

Next, we assessed whether the task dependence of ICMS reflected a modulation of the direct input from S1. To this end, we examined the strength of the pulse-locked response across tasks and found it to be highly consistent. In contrast, neurons whose ICMS-evoked activity was not locked to the ICMS pulses exhibited complex and largely idiosyncratic task dependence (Figure 6C). We conclude that the task dependence of ICMS-evoked activity in M1 does not reflect a change in the direct input from S1 but rather a change in the impact of this input on M1.

### ICMS-evoked M1 activity contaminates motor decoding

Finally, we examined whether the ICMS-evoked M1 activity had an impact on the ability to decode motor intent. To this end, we trained an Optimal Linear Estimator decoder^7^ to control three translational degrees of freedom, with which participant C1 controlled a virtual arm to reach to an object, grasp it, and transport it to a new location. In these experiments, the grasp was automatically triggered once the hand reached the object’s location, to decouple the ICMS from the grasp kinematics, thereby ensuring that the ICMS was identical across grasps. During object contact, ICMS amplitude was constant, at an amplitude corresponding to the grasp force required to maintain object grasp (i.e., evoking a moderately strong tactile sensation), as is typically done^8^. The addition of linear stimulation led to significantly more failed trials, in which the participant was unable to complete the transport within the 10-second window (38% vs. 12%, with and without stimulation, p < 0.001, chi-squared test; Figure 7A). These failures were primarily due to movements being less efficient, with both path lengths and completion times during transport being longer with stimulation (4.6 m vs. 1.4 m, 8.4 s vs 2.6 s, both p < 0.001, two-sample

### Biomimetic sensory feedback rescues decoder performance

Importantly, because ICMS-evoked activity in M1 is task dependent, its influence on a decoder cannot be easily eliminated. However, we reasoned that reducing the amount of ICMS would reduce its deleterious effects. With peripheral nerve interfaces, biomimetic somatosensory feedback – characterized by high-amplitude phasic stimulation at the onset and offset of contact, and far weaker stimulation during maintained contact^9,10^ – has been shown to elicit more natural and intuitive sensations ^11,12^. In the present context, we reasoned that this feedback might offer the additional benefit of reducing the total amount of stimulation, thereby improving decoding. To test this possibility, we had the participant perform the reach, grasp, and transport task but provided information about grasp force using biomimetic ICMS-based feedback. With biomimetic feedback, the onset and offset transient amplitudes were higher than the highest amplitude used in the linear trains but sustained stimulation was weaker. Nevertheless, with this feedback, the participant transported objects with a third as many failures compared with linear stimulation (12% vs. 38%, p < 0.001, chi-squared test; Figure 7A). This improvement was due to shorter, quicker movements during the transport phase (1.4 m vs. 4.6 m, 2.6 s vs 8.4 s, both p < 0.001, two-sample Kolmogorov-Smirnov test; Figure 7B; Supplementary Figure 13). In fact, performance with the biomimetic feedback was nearly identical to that with no stimulation (12% vs. 12% failure rate, p = 0.87, chi-squared test; mean path length and mean trial duration were numerically identical for both conditions at 1.4 m and 2.6 s, respectively; Figure 7; Supplementary Figure 13). Note that ICMS-feedback did not have any beneficial effects on performance here because grasping was automated and therefore did not require or allow for any online correction.

## Discussion

We show that ICMS of S1 evokes widespread activity in M1. Some of this activity takes the form of short-latency responses to ICMS that are phase-locked to individual ICMS pulses. Most of the ICMS-driven activation in M1 is not pulse-locked, however, and seems to reflect an indirect effect of S1 input. The spatial pattern of evoked activity in M1 depends systematically on the location of the S1 stimulating electrode: an M1 channel is susceptible to being modulated by an S1 channel to the extent that they both encode a matching part of the hand. The ICMS-dependent M1 modulation is task dependent, but only for neurons that appear to be indirectly activated by ICMS. In other words, the signals that are directly transmitted from S1 to M1 are consistent across tasks, but their indirect effects are not. Finally, ICMS-evoked M1 activity is relevant to prosthetics as it disrupts the ability to decode motor intent. However, this disruption can be minimized with a more biomimetic form of somatosensory feedback, which emphasizes the transient phases of object contact (onset, offset) and minimizes sustained ICMS.

### S1 sends signals directly to M1

In both humans and macaques, Brodmann’s area 1 and M1 have been shown to be connected anatomically^3,4^. In macaque monkeys, tracer injections in area 1 reveal reciprocal connections with M1^3^, albeit sparse ones. In humans, probabilistic diffusion tractography reveals strong connections between area 1 and M1^4^. Microstimulation of human somatosensory cortex with either surface or penetrating electrodes has been shown to evoke field potentials in motor cortex^5,6^, revealing a functional correlate to the anatomical findings. However, neither the time course of these signals nor their spatial specificity could be gleaned from these measurements of aggregate neuronal activity in M1. While short latency ICMS-evoked responses have been found across sensorimotor cortex in other organisms ^13–17^, the present report is the first to document systematic signaling between somatosensory and motor cortices of humans at the cellular level. Some of the short-latency, low-jitter M1 responses to ICMS in S1 may reflect antidromic activation, but the latency, jitter, and spiking probabilities of the pulse-locked responses were smoothly distributed over a range, offering no hint of a separation between two classes of activation (antidromic vs. orthodromic)(Supplementary Figure 6).

### Body maps in somatosensory and motor cortex are linked

We found that the functional connectivity between S1 and M1 is systematically patterned: neighboring electrodes in S1 produce similar spatial patterns of activation in M1. Moreover, this patterning follows somatotopic maps in S1 and M1: a given channel in S1 is liable to activate a given channel in M1 to the extent that these encode overlapping parts of the hand. The somatotopic patterning in M1 seems at odds with the observations that individual M1 neurons encode movements of joints distributed over the entire hand^18–20^, resulting in a coarse somatotopic organization. Nonetheless, we did observe a coarse somatotopic progression over the sampled cortex, even within the M1 hand representation. The somatotopic organization of the S1-M1 connectivity is consistent with the interpretation that sensory feedback from a given digit preferentially informs the ongoing motor control of that digit. Note, however, that we were also able to decode reaching movements from the putative hand representation in M1, arguing this somatotopic organization is not absolute, consistent with prior findings ^21^. Interestingly, while ICMS delivered to hand S1 disrupts decoding of reaching movements from hand M1 (in participant C1), ICMS delivered to hand S1, in a previous study, seems to have had little impact on the ability to decode reaching movements from arrays located more medially in M1, presumably farther from the hand representation (in participant P2^8^). The somatotopically linked connectivity observed here may underlie the observation in macaques that M1 neurons receive tactile input on the associated hand segment^22^. Analysis of the nature of these signals during natural manual interactions in intact humans and monkeys will shed further light on the functional role of this cortico-cortical signaling.

### ICMS-evoked activity in M1 is relevant to neuroprosthetic development

ICMS of S1 has been shown to elicit vivid sensations that are experienced on the hand^23–27^. These sensations can be used to provide tactile feedback about object interactions and have been shown to improve the functionality of a brain-controlled robotic hand^8^. In the one demonstration of the benefits of somatosensory feedback on object manipulation, the participant’s motor arrays were located in the proximal limb representation of M1 and were only weakly impacted by ICMS to S1 (P2 in this study, see Supplementary Figure 1). When M1 and S1 arrays are both in the respective hand representations, ICMS has a deleterious effect on decoding, thereby counteracting – at least in part – the benefits of sensation. The fact that ICMS-induced activity in M1 is dependent on behavior on a subset of channels – those for which the influence of stimulation is indirect (Figure 6C) – implies that mitigating the impact of ICMS on decoding will be challenging. Indeed, training a decoder based on combined observation and stimulation will work only (1) if the decoder is trained on tasks that span the space of possible behaviors and (2) if the subspace of ICMS-evoked activity in M1 is largely non-overlapping with that involved in motor control. The first condition will be impossible to meet given realistic time constraints, and we have evidence that the second condition is not met (Supplementary Figure 14). However, the impact of ICMS on decoding was largely eliminated by implementing phasic biomimetic feedback, designed to mimic natural cutaneous responses in cortex. In experiments with electrical interfaces with the peripheral nerve, biomimetic sensory feedback has been shown to be more intuitive and naturalistic^11,12,28^. Here, we show that biomimetic stimulation may also alleviate the disruptive effect of ICMS on decoding performance for brain-controlled bionic hands.

## Conclusion

ICMS in S1 reveals strong signaling from S1 to M1 that is patterned such that S1 neurons with projected fields on one hand region preferentially activate M1 neurons that are implicated in moving that hand region. While the direct connection between S1 and M1 is fixed, the overall impact of ICMS to S1 on M1 activity is task dependent. This channel of communication between S1 and M1 disrupts the decoding of motor intent from M1 signals, but this disruption can be minimized using biomimetic feedback.

## Methods

### Participants

The three participants, part of a multi-site clinical trial (NCT01894802), provided informed consent prior to any experimental procedures. Participant C1 (male), 57 years old at the time of implant, presented with a C4-level ASIA D spinal cord injury (SCI) that occurred 35 years prior. He had no spared control of the intrinsic or extrinsic muscles of the right hand but retained the ability to move his arm with noted weakness in many upper limb muscles. Filament tests revealed spared deep sensation but diminished light touch in the right hand (detection thresholds ranged from 0.6 to 2.0 g across digit tips). Participant P2 (male), 28 years old at the time of implant, presented with a C5 motor/C6 sensory ASIA B SCI that occurred 10 years prior. He had no spared control of the intrinsic or extrinsic muscles of the right hand but had limited control of wrist flexion and extension. Proximal limb control at the shoulder was intact, as was elbow flexion. However, he had no voluntary control of elbow extension. He was insensate in the ulnar region of the hand (digits 3-5) on both the palmar and volar surfaces but retained both diminished light touch and deep sensation on the radial side (digits 1-2) (thresholds were 1.4 g to 8 g on the thumb and index, respectively, and 180 g on the middle finger). Participant P3 (male), 28 years old at the time of implant, presented with a C6 ASIA B SCI that occurred 12 years prior. He had no functional control of the intrinsic or extrinsic muscles of the right hand but retained the ability to move his arm with noted weakness in many upper limb muscles. He was insensate in the ulnar region of the hand on both the palmar and volar surfaces but retained diminished light touch and deep sensation on the radial side (thresholds were 0.07 g and 1.6 g on the thumb and index and 8 g on the middle finger).

### Array implantation

We implanted four Neuroport microelectrode arrays (Blackrock Neurotech, Salt Lake City, UT, USA) in the left hemisphere of each participant. Two of the arrays, implanted in somatosensory cortex (S1), were 2.4 mm x 4 mm, each with sixty 1.5-mm electrode shanks wired in a checkerboard pattern such that 32 electrodes could be stimulated. The other two arrays, implanted in motor cortex (M1), were 4 mm x 4 mm with one hundred 1.5-mm electrode shanks, 96 (participants C1 and P3) or 88 (participant P2) of which were wired (active). Four inactive shanks were located at the corners of all arrays (with an additional 8 for participant P2). In P2, the motor cortex arrays were metalized with platinum while the somatosensory arrays with coated in sputtered iridium oxide. In participants C1 and P3, all electrodes were coated with sputtered iridium oxide. Most of the electrodes (74/96) on the medial array of participant C1 were too noisy to yield useful data and deactivated. Each participant had two percutaneous connectors placed on their skull, with each connected to one sensory and one motor array. We targeted array placement during surgery using functional neuroimaging of the participants attempting to make movements of the hand and arm, and imagining feeling sensations on their fingertips^23^, within the constraints of anatomical features such as blood vessels and cortical topography (Figure 1A and Supplementary Figure 1). Array locations, shown in Figure 1A and Supplementary Figure 1 on structural MRI models of each participant’s brain, were confirmed using intraoperative photographs after insertion.

### Neural stimulation

Stimulation was delivered using a CereStim microstimulator (Blackrock Neurotech, Salt Lake City, UT, USA). Stimulation pulses were cathodal first, current controlled, and charge balanced, over a range that has been previously deemed safe^29^. Each pulse consisted of a 200-μs long cathodal phase, then a 100-μs interphase period followed by a 400-μs long anodal phase at half the cathodal amplitude. Stimulation pulses could be presented at up to 300 Hz. Further details on selection of stimulation parameters can be found in Flesher et. al. 2016^23^.

### Neural recordings

Neural signals in M1 were recorded at 30 kHz using the NeuroPort system (Blackrock Neurotech, Salt Lake City, UT, USA). Each stimulation pulse triggered a 1.6-ms sample-and-hold circuit in the preamplifier to avoid saturating the amplifiers and to minimize transient-induced ringing in the filtered data. The data were high-pass filtered with a 1st order 750 Hz filter^30^. Whenever the signal crossed a threshold (−4.25 RMS, set at the start of each recording session), a spiking event was recorded and a snippet of the waveform was saved. Spikes were binned in 20-ms bins for decoding. To confirm that the observed effects reflect neural activity and not an electrical artifact, we sorted units offline using Plexon Offline Sorter and repeated many of the analyses described below on isolated single units.

### Stimulation protocol – passive condition

To study the effects of stimulation on M1 activity, we stimulated through each S1 channel a minimum of 15 times at 60 μA and 100 Hz in 1 second trains. Electrode order within each array was shuffled and stimulation was interleaved across arrays. The interval between pulse trains was 3 seconds in participant C1 and a random duration between 3 to 4 seconds in participants P2 and P3 to counteract any anticipatory effects to the stimulation.

### Gauging the strength of ICMS-driven activity in motor cortex

To understand the effects of ICMS in S1 on activity in M1, we compared the fluctuations in firing during baseline to those during the stimulation interval. For each motor channel, we sampled the difference in firing rate between two consecutive 1-second intervals during the intertrial periods, computed the mean, and repeated this process 1000 times to generate a null distribution of baseline fluctuations over the course of a recording session. For each stimulating channel, we calculated the change in firing rate between a 1-second interval preceding the stimulation train and the firing rate during the stimulation train itself, which gauged the effect of stimulation on each motor channel. For these analyses, we extended the blanking window to 2 ms to eliminate any potential electrical artifacts that were not excluded during the initial blanking window. We simulated this blanking in the baseline response to generate the null distribution. Motor channels were considered to be modulated by stimulation if their average change in firing rate during stimulation was significantly different from the null distribution (p < 0.001). To gauge the sign and magnitude of the effect of stimulation on a motor channel, we expressed the change in firing rate during stimulation for each motor/stimulation channel pair as a z-score based on the null distribution for that motor channel. Positive modulation values indicate an excitatory effect while negative modulation values indicate an inhibitory effect.

### Gauging the timing of ICMS-driven activity in motor cortex

To determine if motor units were phase locked to the stimulation pulses, we computed the pulse triggered average (PTA). Specifically, we binned the spikes evoked during each inter-pulse interval into 0.5 ms bins and computed the probability of spiking in each bin (i.e., the proportion of times a pulse evoked a spike in that bin). To assess whether there was a significant peak in the PTA, indicating a pulse-locked response, we first identified the time at which the probability of a spike occurring was highest and averaged the spiking probability across it and the two adjoining time bins. We computed the median probability of a spike occurring across all bins in the inter-pulse interval, to quantify the component of the response that was not pulse-locked. We computed the difference between these two values to create a phase-locking index. We sampled 20% of the PTAs for each motor and stimulation channel pair, shuffled the spike times, thus obtaining PTAs that were matched in spike count, and computed the same phase-locking index above for PTAs generated from the shuffled data. We repeated this shuffling procedure 5000 times to create a null distribution of pulse-locking indices. PTAs were considered to be significantly pulse-locked if the index was greater than that 99% of those obtained by chance (i.e., p < 0.01). We also estimated the latency and jitter of significantly pulse-locked responses. To this end, we randomly sampled 20% of the inter-pulse intervals and computed the PTA for this sample. We then identified the bin with the maximum spiking probability thus determining its latency. We repeated this procedure 5000 times to get a distribution of latencies, the mean and variance of which were the latency and jitter estimate for that stimulation/recording pair.

### Quantifying somatotopically mapped connectivity

We sought to determine whether motor channels that encode information about specific digit movements also respond to stimulation in somatosensory cortex that evokes a touch sensation on the same digit. To this end, participants performed a digit (attempted) movement task. On each trial, a digit was cued and the participant attempted to flex then extend the digit before the next digit was cued. Participant C1 was cued by the name of the digit being spoken, then attempted to move his own paralyzed digit in synchrony with a virtual reality model (MuJoCo, DeepMind Technologies, London, UK) that completed the same instructed movement. He completed 125 trials of this task in one session. Participants P2 and P3 were cued by watching a set of 5 colored circles displayed on a monitor in front of them. The circles were arranged to mimic the distribution of digit tips resting open on a table or keyboard. When a circle was filled by a gray dot the participant would attempt to flex the corresponding digit until the gray dot disappeared. Following a chime, he then attempted to extend the same digit. Each participant completed 50 trials of this task.

<u>Motor maps</u>. To generate a map of digit selectivity across M1, we first computed the mean peri-event time histogram (in 20-ms bins) for each motor channel across a two second period centered on the start of movement for each digit flexion. From these, we then identified, for each motor channel, the response window during which the difference between the maximum response and the minimum response (each corresponding to flexion of different digits) was maximal. We used different time windows for different M1 channels because some units were most strongly active during preparation and others during movement. The modulation value for each digit was then calculated by subtracting the mean firing rate across all digits from the average firing rate for one digit, and then dividing by the mean firing rate across all digits. Plotting this modulation value across all channels for one digit provides a map of selective activation for that digit.

To generate <u>sensory projection maps</u>, we first computed the modulation value for each motor channel when stimulation was delivered through a stimulation channel that evoked a perceived sensation on a given digit. For example, we computed the modulation value for each motor channel when all the stimulation channels with projected fields on the thumb were stimulated. We then averaged these modulation value to obtain the thumb projection map. We repeated this procedure for all the digits (excluding the little finger for participant C1, because he never reported a sensation there).

We compared these two maps – motor and sensory projection – by first convolving them map with a 2D Gaussian with a standard deviation equal to the spacing between two adjacent electrodes, to emphasize large scale spatial patterns of activation and obscure small irregularities. We then computed the covariance between the motor and sensory projection maps (excluding the little finger for participant C1) and compared them to the covariance across 1000 maps in which the electrode locations were randomly shuffled within each digit. The actual maps were considered to be significantly similar if the observed covariance was larger than 99% of those derived from the shuffled maps (p < 0.01).

### Assessing the task dependence of ICMS-evoked activity in M1

We sought to determine whether the effects of ICMS to S1 on M1 activity depended on the task. To this end, we had participants C1 and P3 perform two tasks while we delivered ICMS to S1.

In the first task, the <u>squeeze task</u>, the participant squeezed a virtual object and reported the intensity of the ICMS-evoked touch sensation. On each trial the participant attempted to squeeze a virtual object with a medium amount of force, following the trajectory of a virtual hand observed through a VR headset. Upon contact with the object, stimulation was delivered on two electrodes at one of four amplitudes (20, 32, 44 or 56 μA). The hand continued to grasp the object for one second before a release cue appeared. Once the hand released the object, the participant reported the perceived intensity of the stimulation using a scale of their choosing, with the following instructions. If they did not feel the stimulus, they were to ascribe to the sensation a rating of zero. If a stimulus on one trial felt twice as intense as that on another, it was to be ascribed a rating that was twice as high (other such examples were provided). They were encouraged to use decimals or fractions. The main goal of the magnitude estimation component was to keep the participant engaged in the task.

The second task was a <u>grasp and transport task</u>. On each trial, an object appeared at one corner of an invisible cube centered on the starting point of the virtual hand. The participant then attempted to reach to the object, following the movement of the virtual hand. Once there, the participant attempted to grasp the object with medium force. During the grasp, ICMS was delivered at one of four amplitudes, as in the magnitude estimation task. The participant then attempted to bring the object back to the center of the cube and release it there, again following the movements of the virtual limb.

Participant C1 completed 208 trials of each task across two sessions. Participant P3 completed 160 trials of each task in one session.

To confirm that the participant was attending to the grasp and transport task, we classified the intended target during the reach phase of the task. A naïve Bayes classifier was trained using one second of data from all active motor channels (>5 Hz mean firing rate across whole task) starting 400 ms before movement onset. This classifier was tested using leave-one-out cross-validation.

To assess whether the ICMS-evoked M1 activity varied across tasks, we analyzed the firing rates across all motor channels during three distinct phases across the two tasks: The one second period after grasp contact during ‘squeeze’ task; the one second period after grasp during the grasp and transport task, and the first second of the transport phase in the grasp and transport task. In all three of these phases, the ICMS was identical but the movements were different (squeeze/grasp vs. transport) or their context was different (squeeze vs. grasp). We performed a multivariate ANOVA on the firing rates to determine which channels were significantly modulated by changes in task phase, stimulation level, and the interaction of the two. As an index of task dependence, we computed the ratio of the F-statistic for the interaction effect to that for the main effect of stimulation. This value was high to the extent that the interaction effect was strong compared to the main effect. This index was only computed for significantly modulated channels.

To further determine how different the ICMS-evoked M1 activity was across tasks, we trained naïve Bayes classifiers on stimulation amplitude during the squeeze, grasp, and transport phases separately and tested these classifiers on all three conditions. Within-condition accuracy was calculated using leave-one-out cross-validation, while cross condition accuracy was calculated using a decoder calibrated with all available trials.

### Quantifying the impact of ICMS on motor decoding

We sought to determine whether the motor cortical activity evoked by ICMS would disrupt the ability of the participants to control a virtual arm. To this end, we trained a decoder using the methods described previously^7,8^. Briefly, in 3 sessions, participant C1 attempted to make the movements of the virtual hand and arm displayed in his VR headset. On each trial, the virtual hand reached to an object, grasped it, transported it to a new location, and released it. After completing 60 trials, we trained a decoder for three-dimensional translation of the hand using these data. Next, we measured neuronal activity as the participant controlled translation, but with the computer preventing deviations from the path to the target (for an additional 60 trials). A new decoder was then trained from these data, and that decoder was used for the rest of the session. The decoders were trained without stimulation but with blanking applied at 100 Hz during object contact to simulate the neuronal signal available during online decoding with ICMS-based feedback.

Once the decoder was trained, performance was tested under three conditions. In the ‘no stimulation’ condition, the participant performed the same task that was used during training; no stimulation was provided but a 1.6-ms window of neuronal data was blanked at 100 Hz to match the data available during stimulation. In the ‘linear stimulation’ condition, ICMS was delivered on two electrodes, one with projected fields on the thumb and one on the index finger. In the linear condition, the ICMS frequency was 100 Hz and the amplitude was 52 μA. In the ‘biomimetic stimulation’ condition, 100-Hz ICMS was delivered through the same two electrodes but had onset and offset transients of 72 μA for 200 ms with 32 μA during maintained contact. The order of the test blocks was randomized in each session, with each condition used for two sets of ten trials before the next condition was tested. Conditions were alternated three times to obtain a total of 60 trials for each.

If the participant was unable to place the hand at the target location within 10 seconds during either the reach or the transport phase, the trial was terminated and marked as a failure. To determine the causes of failure, we computed the path length during the transport phase (when stimulation was provided and the participant had control of the arm) for every trial, even if the trial failed during that phase. The median path lengths were compared across stimulation conditions using the Wilcoxon rank-sum test to determine significance. In the same way, the completion times for the transport phase were compared, with failed trials ascribed a completion time of 10 seconds (the time at which the trial was terminated).

## Acknowledgments

This work was supported by NINDS grants UH3 NS107714 and R35 NS122333.

## Supplementary Figures

**Supplementary Figure 1.**
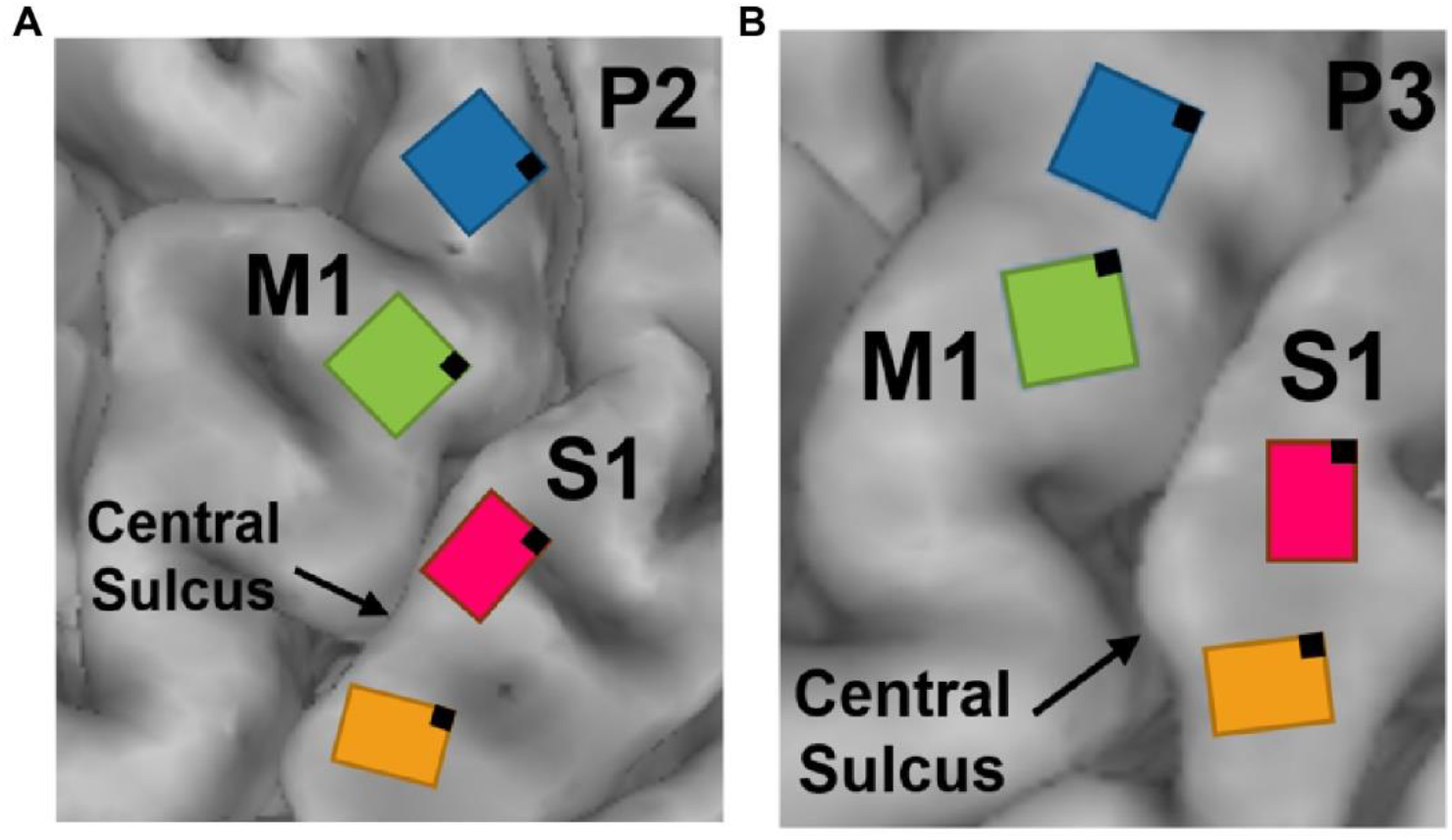
Implant locations for P2 (A) and P3 (B). See Figure 1 for C1.

**Supplementary Figure 2.**
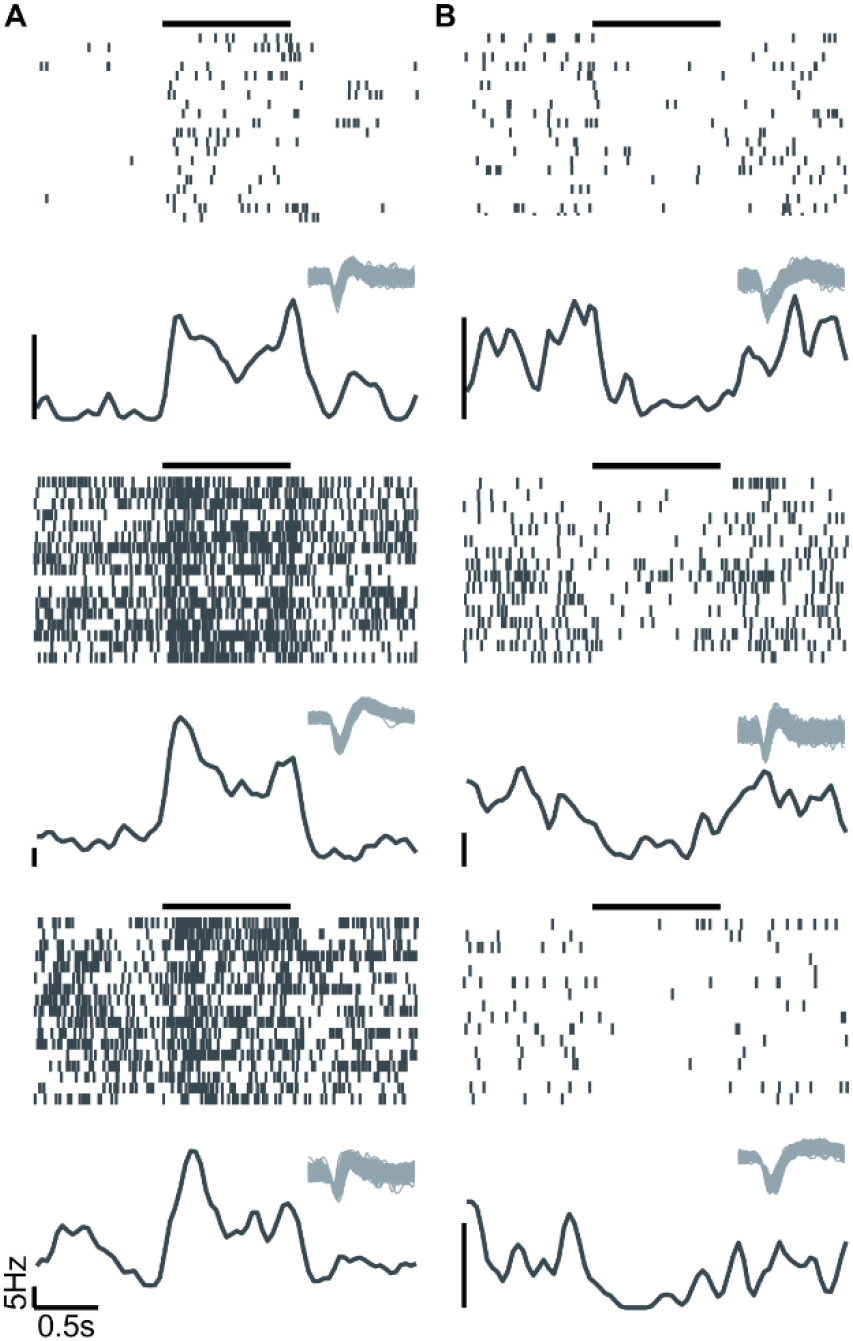
Single unit responses in M1 during ICMS delivered to S1. A) Responses of three example neurons (inset: sorted waveforms) that were excited by stimulation (from top to bottom: C1, P3, P3). B) Responses of three example neurons that were inhibited by stimulation (C1, P2, P3).

**Supplementary Figure 3.**
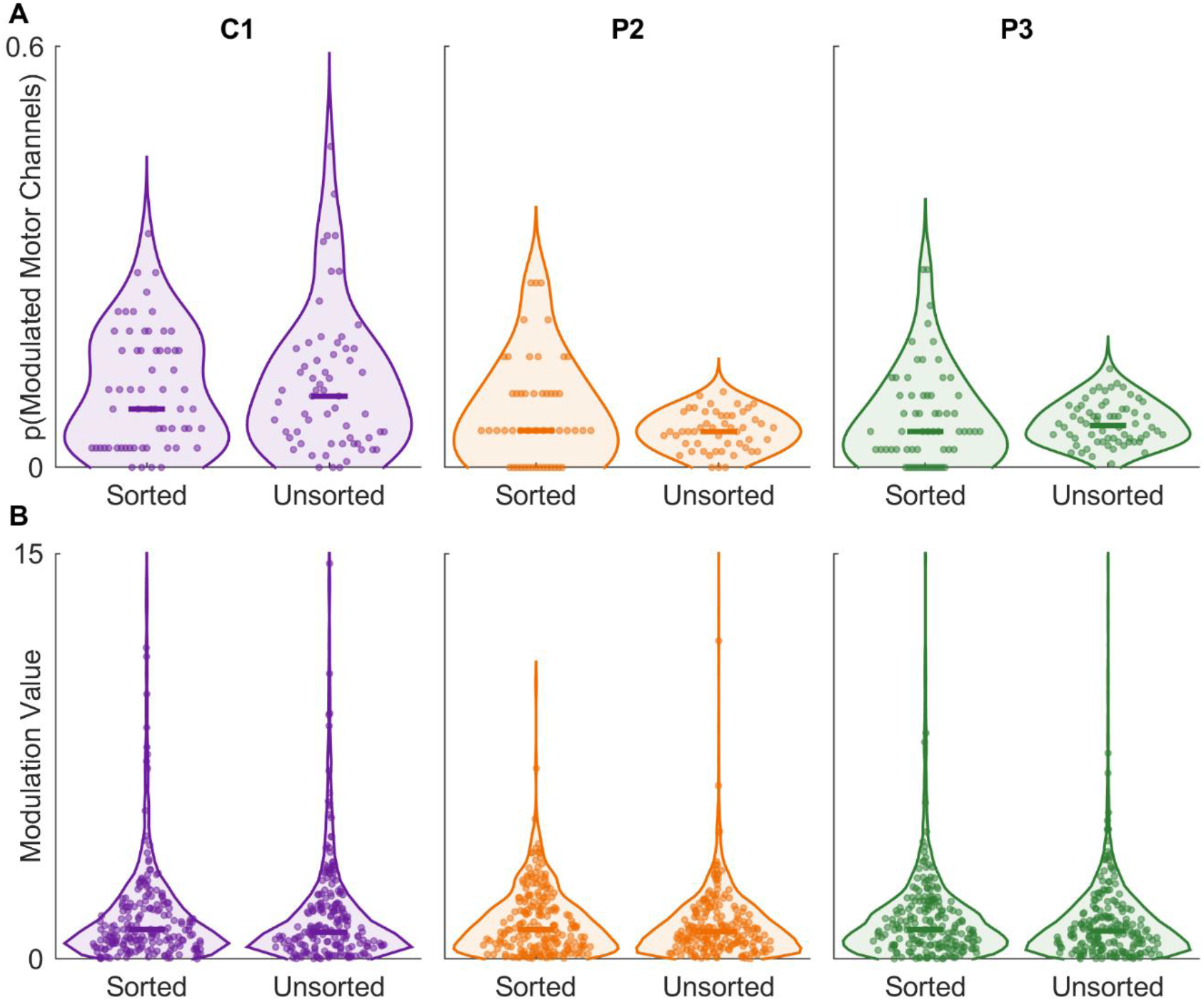
Prevalence of ICMS-evoked activity in motor cortex is similar for sorted and unsorted units (N = 36, 19, and 39 sorted units for participants C1, P2, and P3, respectively). A| Proportion of motor channels significantly modulated by stimulation channels (each dot represents a stimulation channel) for sorted and unsorted units. B| Distribution of absolute modulation values for all pairs of motor and stimulation channels. Unlike in Figure 2B, where modulation values are averaged for each motor channel, the modulation value for each motor-sensory pair is shown separately given the small number of sorted units.

**Supplementary Figure 4.**
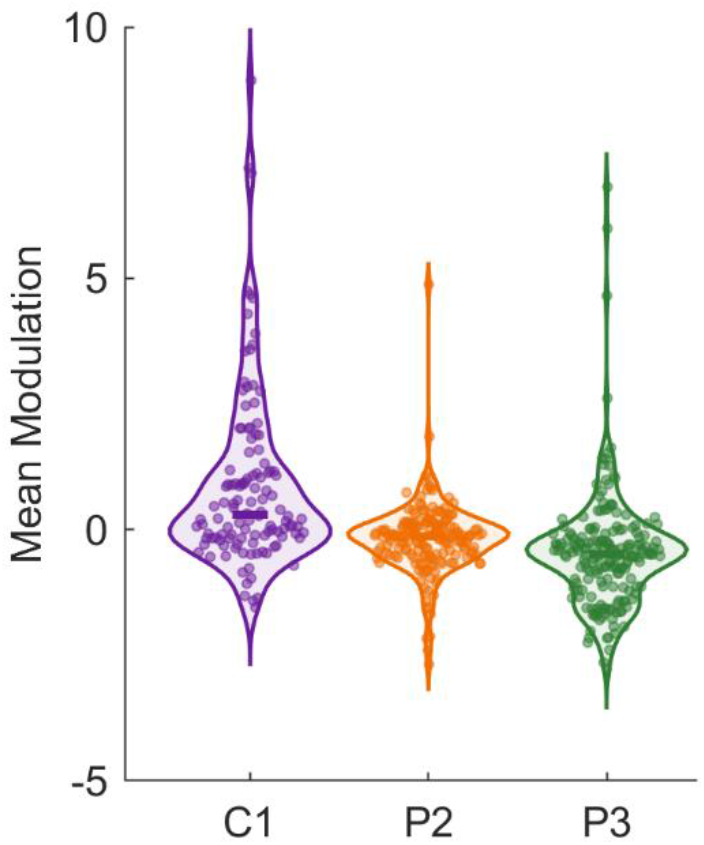
Mean ICMS-driven modulation for each motor channel. This figure shows the raw (signed) modulation values rather than the absolute ones.

**Supplementary Figure 5.**
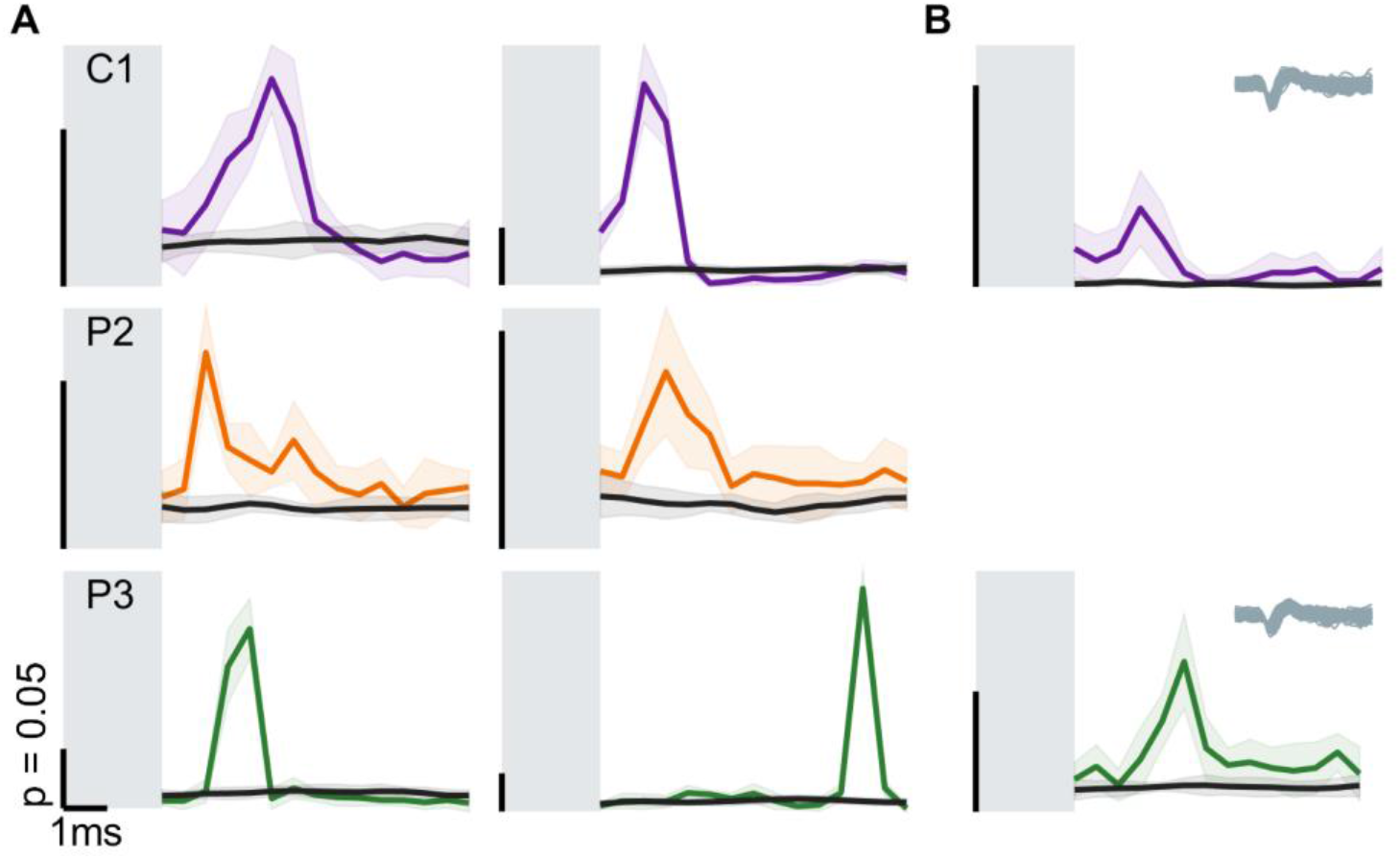
ICMS to S1 evokes short-latency, pulse-locked responses in M1. A| Pulse triggered average (PTA) of the M1 responses evoked ICMS to S1 (colored). As a control, we computed a sham pulse triggered average (at the same pulse frequency) during baseline (black). Temporally precise responses occur at varying latencies across motor channels and participants. Each row shows example PTAs for each participant (C1, P2, P3). The probability of a spike occurring in each 0.5 ms bin is shown on the y-axis. B| Example PTAs for sorted units from participants C1 and P3. There were no sorted units with phase-locking in P2.

**Supplementary Figure 6.**
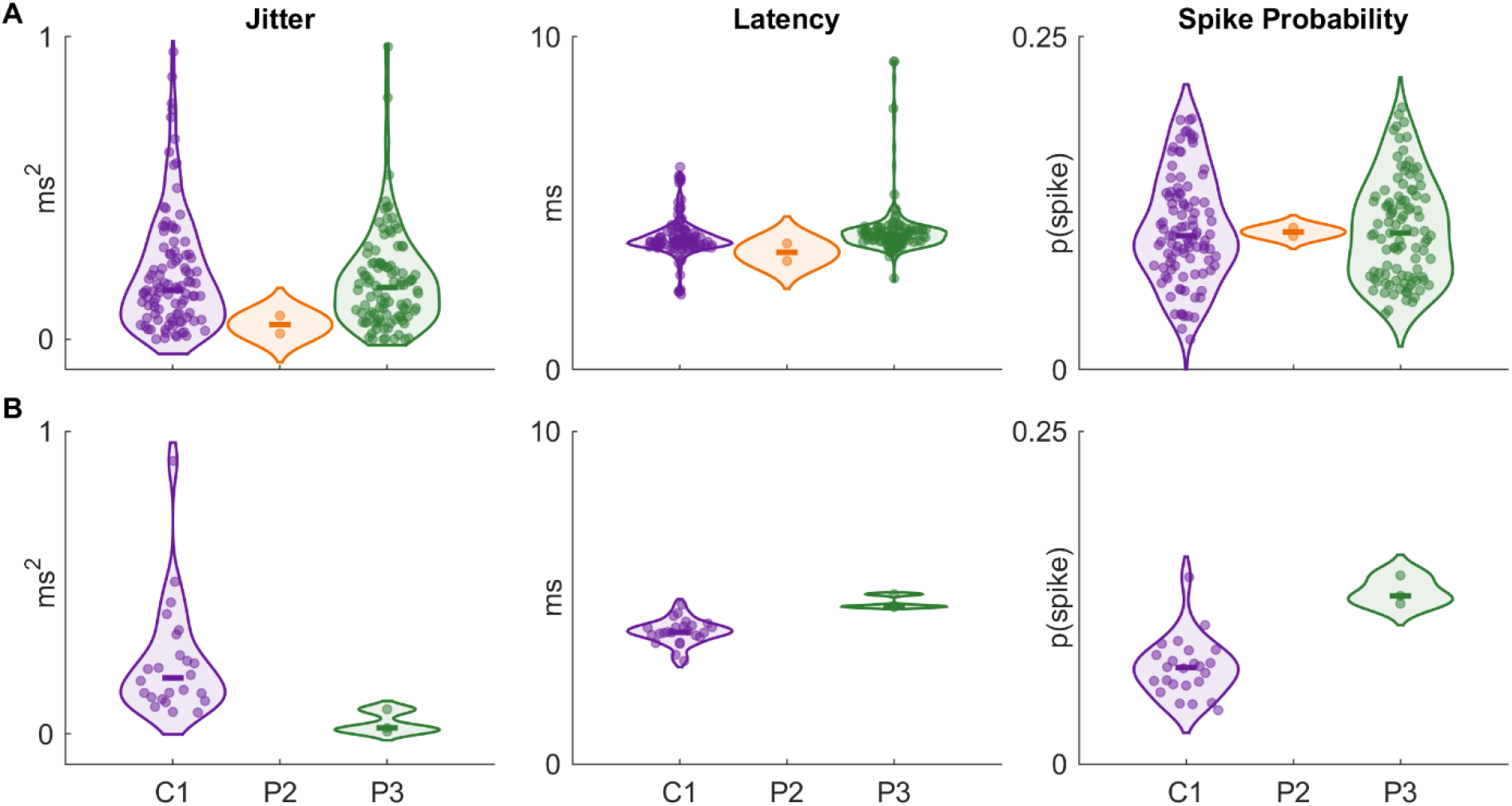
Characteristics of pulse-locked responses. A| Characteristics of pulse-locked responses of unsorted units for each participant. Each dot represents a motor channel-stimulation channel pair. The three metrics – jitter, latency, and spiking probability (the proportion of times a pulse evoked a spike within a 1-ms window centered at the time of highest spiking probability) – were distributed unimodally, precluding classification of pulse-locked responses as reflecting antidromic or orthodromic activation. If spikes with jitter less than 0.1 ms^2^ are considered to reflect antidromic activation (cf. refs. ^31,32^), these responses reflect both antidromic and orthodromic activity, with a far greater prevalence of orthodromic activity. B| The same metrics in sorted units are consistent with responses of unsorted units.

**Supplementary Figure 7.**
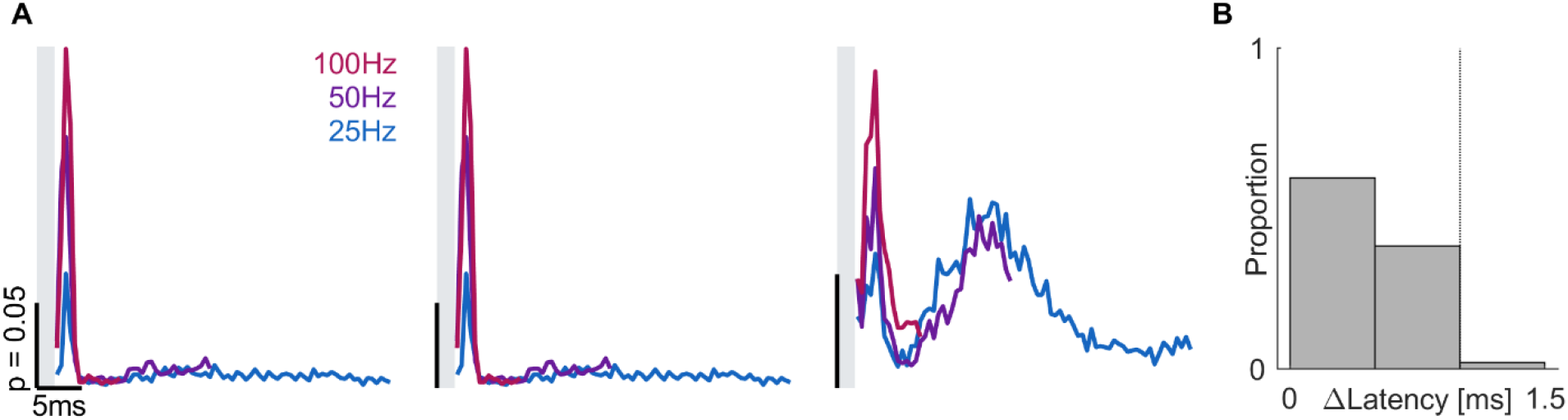
Pulse-triggered average (PTA) of the responses evoked by ICMS at three frequencies. A| Motor channels preserve the initial response latency regardless of stimulation frequency. Some channels demonstrate a secondary, longer latency but lower probability response that is obscured during high frequency stimulation. B| Distribution of the differences in peak latency times across the 3 frequencies. Dotted line indicates the temporal resolution of the analysis. The vast majority of peak latency differences fall below the dotted line, indicating that the time at which the first peak occurs is consistent across frequencies.

**Supplementary Figure 8.**
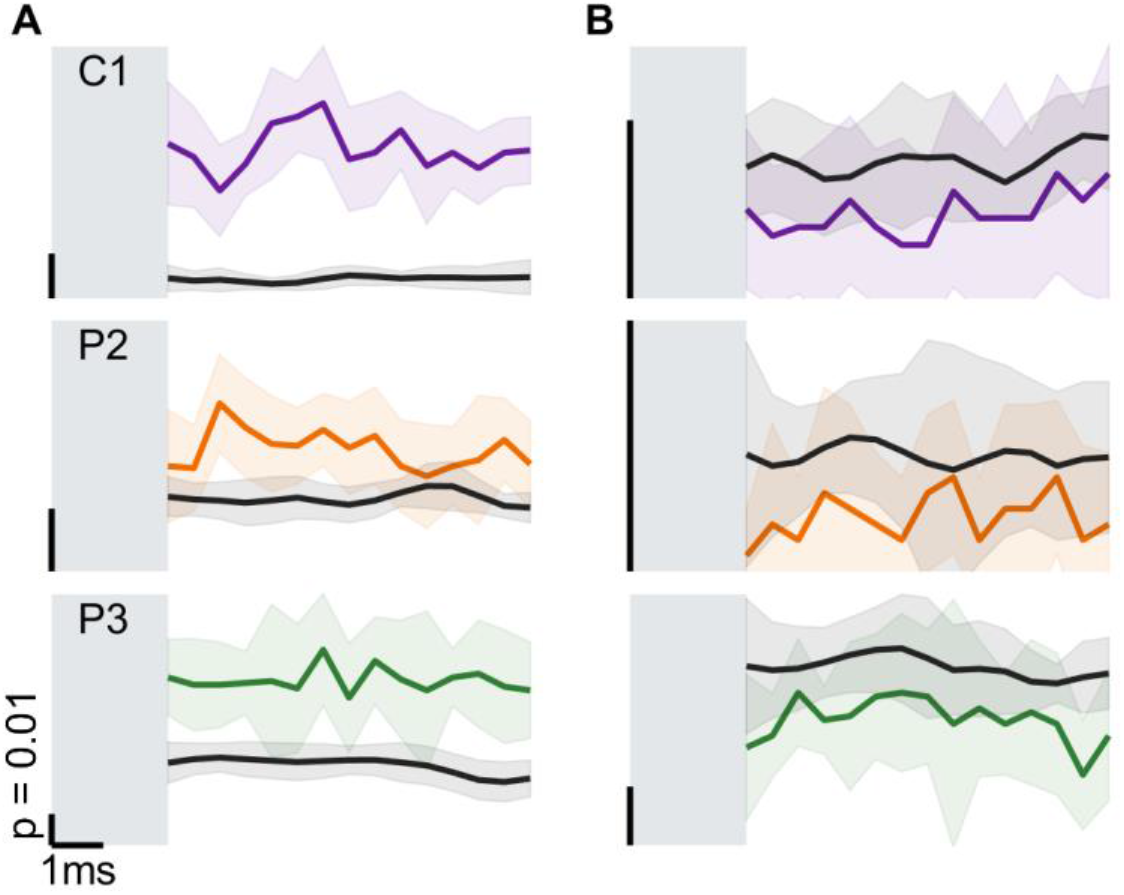
Pulse triggered average (PTA) during ICMS (colored) from channels that were modulated by stimulation but did not exhibit pulse locking in their response. As a control, we computed a sham pulse triggered average (at the same pulse frequency) during baseline (black). The gray area indicates time during which recording was blanked to eliminate the stimulation artifact. The y-axis denotes the probability of a spike occurring in each bin. A| Channels that were significantly excited by stimulation. B| Channels that were significantly inhibited by stimulation. Each row shows example PTAs for each participant (C1, P2, P3).

**Supplementary Figure 9.**
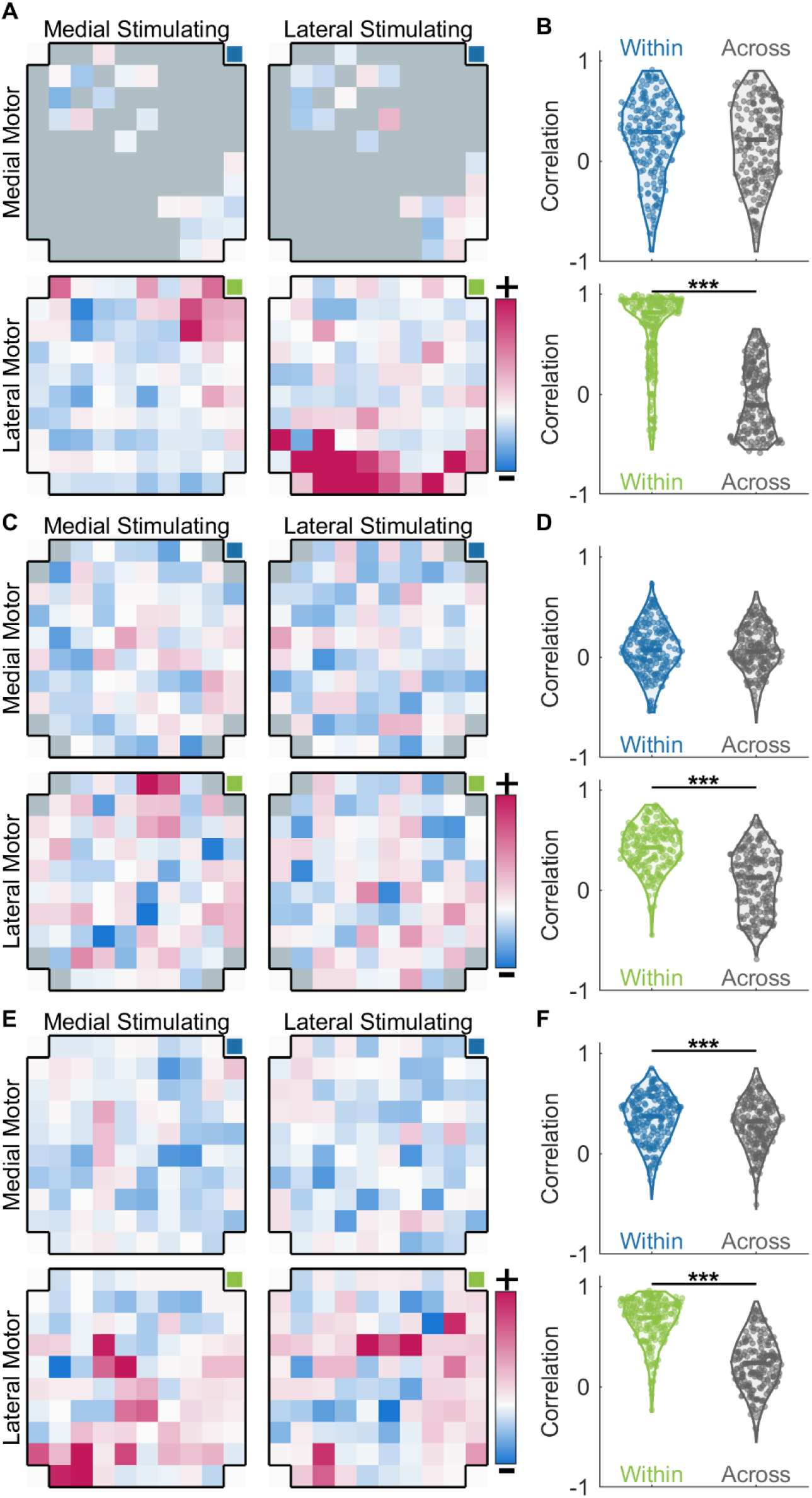
Spatial patterning of ICMS-evoked M1 activation. A| Stimulation through a channel on the medial somatosensory array and lateral sensory array for C1. Adjacent channels are separated by 400 μm. Blue and green squares indicate the orientation of the arrays on cortex (Figure 1A). Grey squares denote inactive motor channels. B| Response correlation between stimulation channels belonging to the same stimulation array (within array) and different arrays (across array) for C1. Asterisks indicate significance (p < 0.001 rank-sum test). C| Same as A for P2. D| Same as B for P2. E| Same as A and C for P3. F| Same as B and D for P3. The ICMS-evoked M1 activity is patterned in C1 and P3 but not in P2.

**Supplementary Figure 10.**
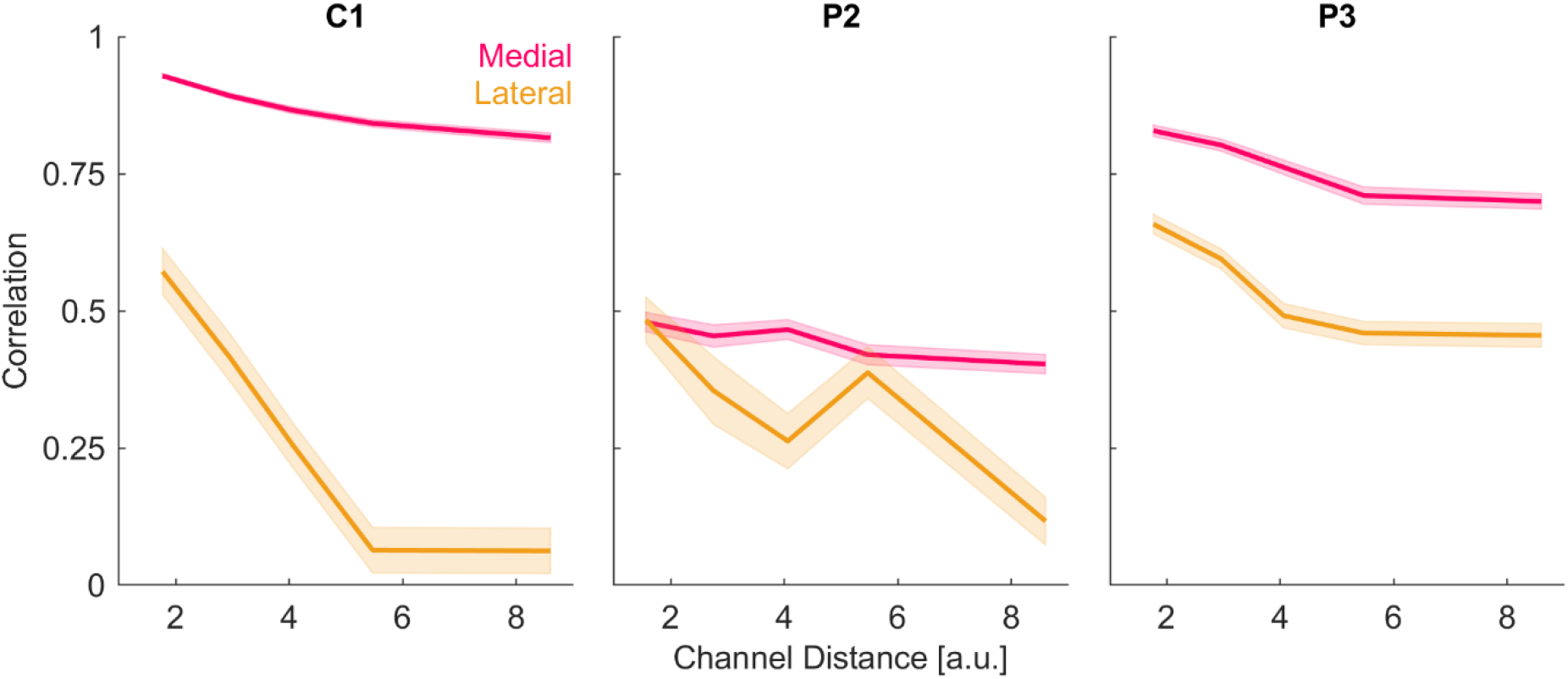
Correlation between the spatial pattern evoked in the lateral motor array as a function of distance between two stimulating electrodes in the medial (pink) and lateral (orange) sensory arrays. The spatial pattern of activation evoked by two electrodes tends to be more similar when the two electrodes are nearby. Correlations were lower for participant P2 overall.

**Supplementary Figure 11.**
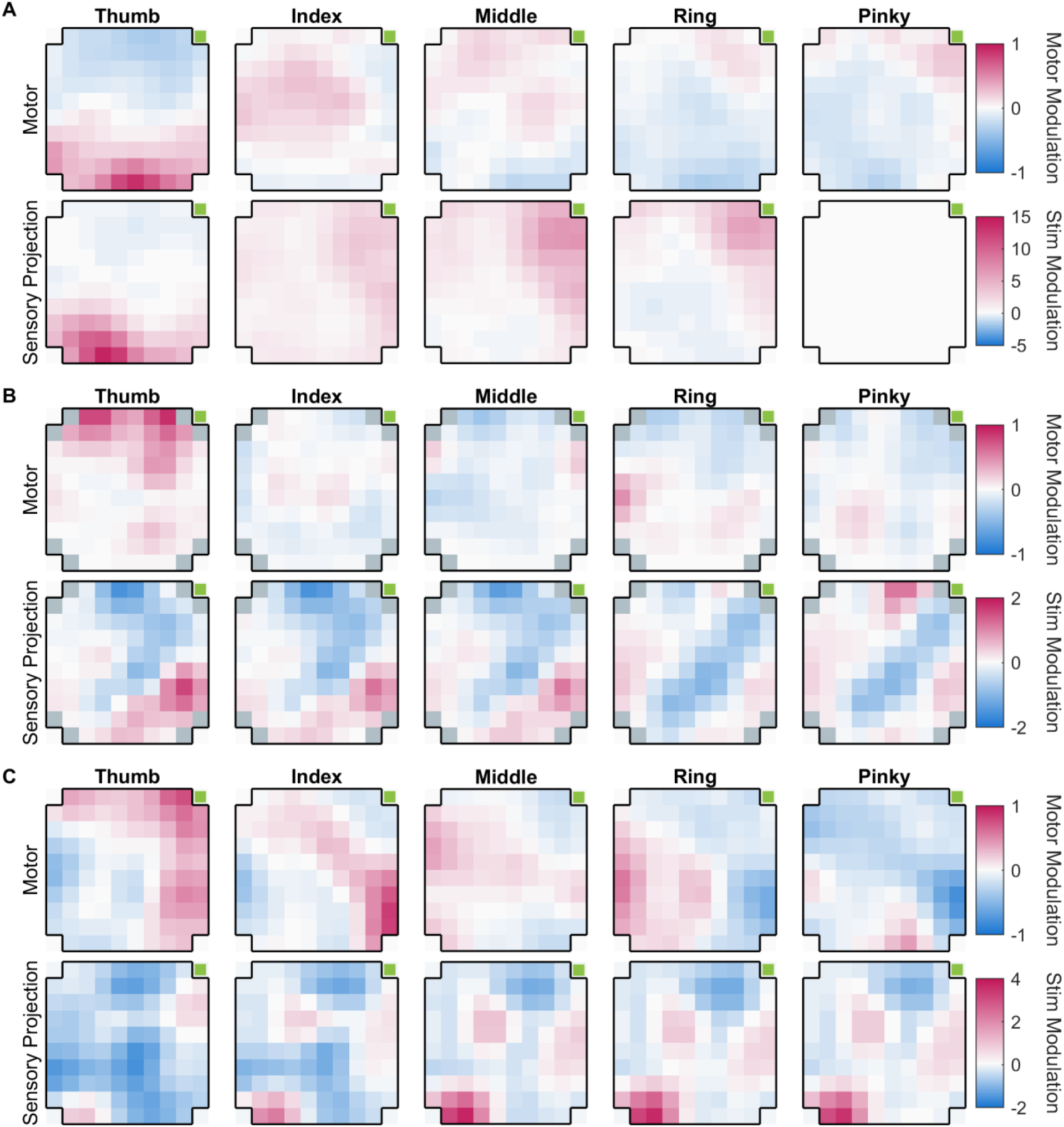
Motor and projection maps for all digits in participants C1, P2, and P3. A | Maps for participant C1. Top: Activation of each motor channel on the lateral motor array for all five digits. Bottom: Average activity evoked by stimulation on channels with projected fields on all five digits (no electrodes had projected fields in the pinky for C1). The green square denotes the medial-posterior corner of the array (see Figure 1A). B| Same as A but for P2, showing the lateral motor array. The green square denotes the medial-posterior corner of the array (see Supplementary Figure 1B). Grey squares denote inactive motor channels. C| Same as A and B but for P3, showing the lateral motor array. The green square denotes the medial-posterior corner of the array (see Supplementary Figure 1C).

**Supplementary Figure 12.**
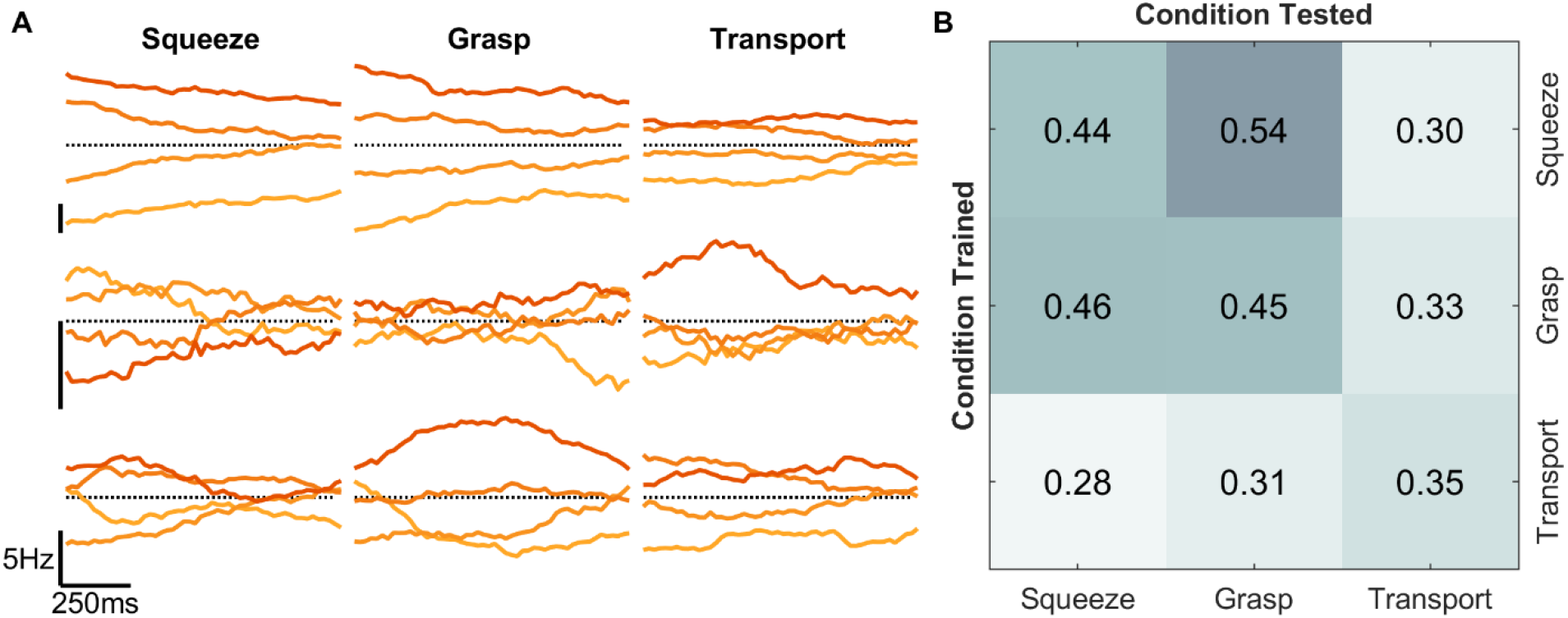
Behavioral modulation of stimulation response in P3. A| Three example motor channels exhibit different responses to four levels of ICMS across three motor conditions (squeeze, grasp, transport). B| Stimulation amplitude classifier performance. Classifiers were trained on one of the three conditions and tested on each condition (with cross-validation for within-condition classification).

**Supplementary Figure 13.**
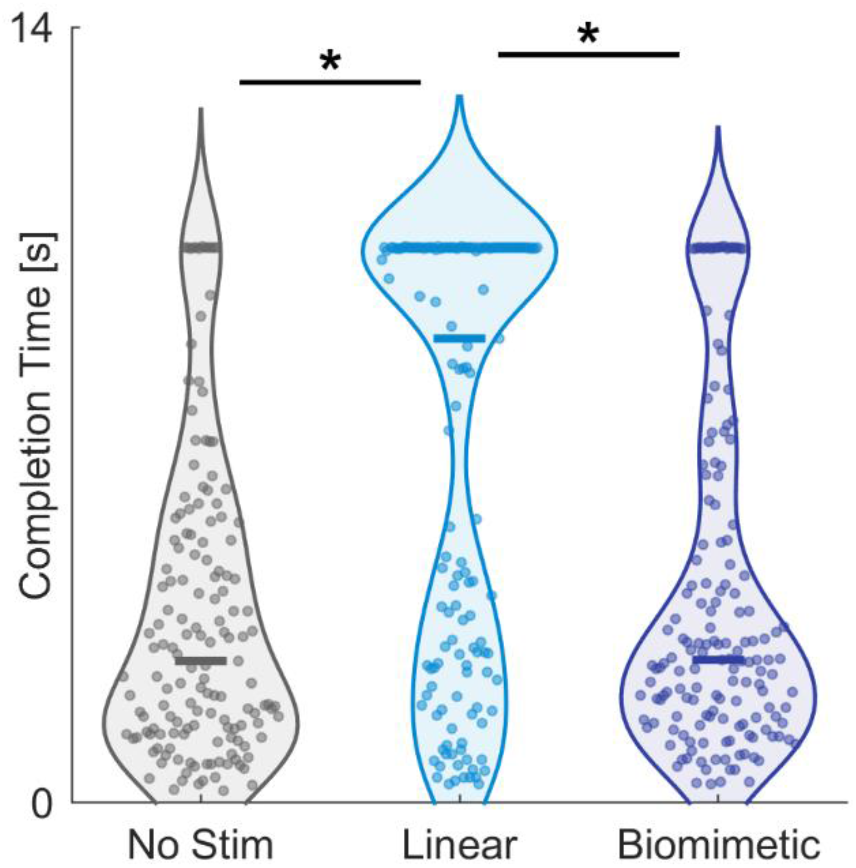
Decoder performance with and without sensory feedback: Transport time. Distributions of the time to complete the transport of an object with no stimulation, linear stimulation, and biomimetic stimulation. Failed trails are denoted by a point at 10 seconds. No stimulation and biomimetic stimulation enabled quicker movements than linear stimulation (p < 0.001, K-S Test). Horizontal bar denotes the median.

**Supplementary Figure 14.**
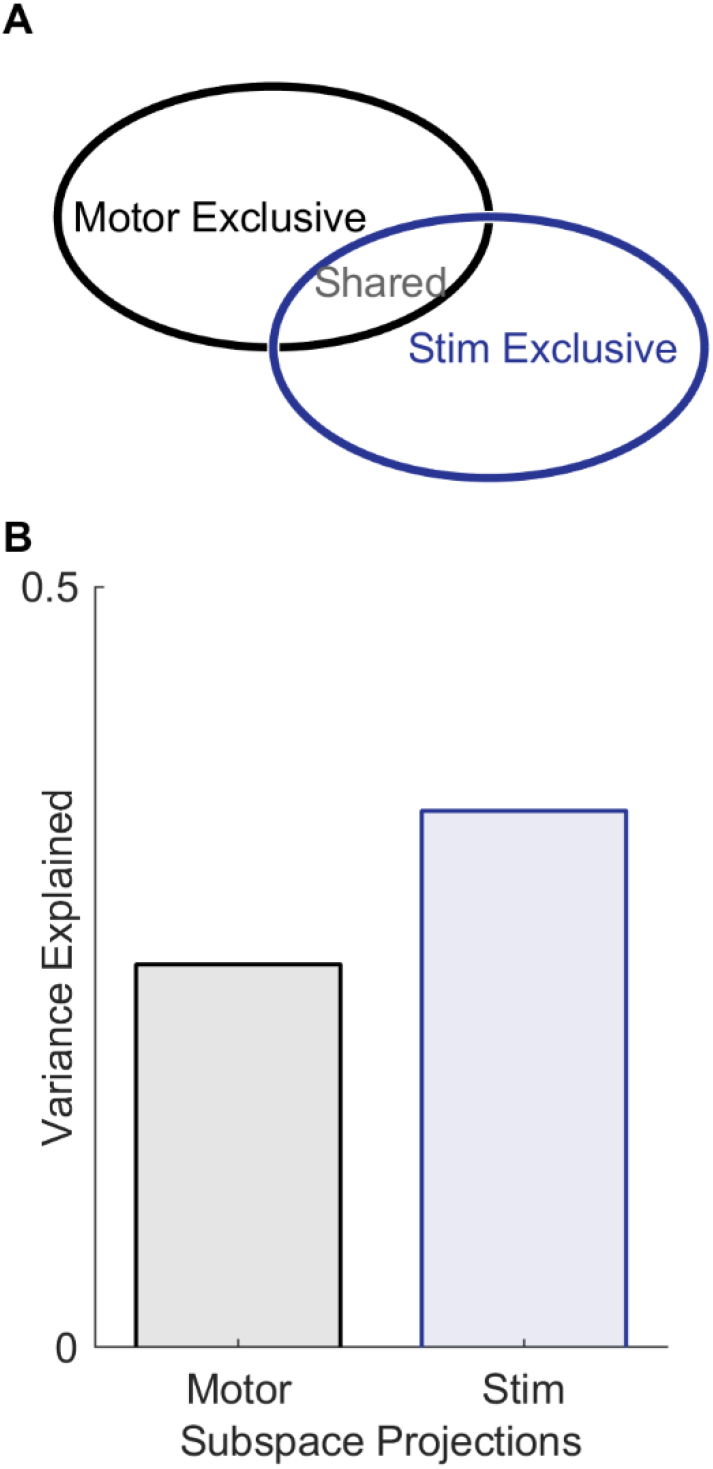
To quantify overlap between motor and stimulation subspaces, we ran two tasks. In one, we instructed the participant to attempt to grasp a virtual object at one of 4 force levels, hold it for 1 second, then release it. No stimulation was delivered during the task. In the second task, we delivered stimulation trains that were identical in duration and shape to the grasp profiles in the first task. The participant was blinded to the level of stimulation and was instructed to report the magnitude of stimulation to maintain engagement in the task. By comparing the M1 activity in the two conditions, we can extract subspaces in M1 population activity that is exclusive to the motor task (related to volitionally moving the hand) or to stimulation, as well as the subspace shared by the two tasks. A| Three subspaces were extracted: One that contains variance of the motor task, one that contains variance of the stimulation task, and one that contains the variance that is common to the two tasks (using methods from ref. ^33^). B| The shared subspace captures significant variance of both motor and stimulation tasks.

